# Rab4 and Rab11 GTPases cooperate to reinforce adherens junctions at the leading edge to promote rapid embryonic wound healing

**DOI:** 10.64898/2026.07.15.738337

**Authors:** Brianna D. O’Donnell, Sofia Mendez-Lopez, Kate D. MacQuarrie, Rodrigo Fernandez-Gonzalez, Katheryn E. Rothenberg

## Abstract

Collective cell movements drive the formation and repair of tissues and contribute to the spread of metastatic disease. Cells must remodel their cell-cell adhesions and actomyosin cytoskeleton to enable migration, but the mechanisms that drive these molecular rearrangements are unclear. We used wound healing in the *Drosophila* embryonic epidermis to investigate these mechanisms. Upon wounding, a supracellular cable composed of actin and myosin assembles around the wound. In parallel, adherens junction proteins, including E-cadherin, are depleted from the wound edge via endocytosis and accumulate at former tricellular junctions around the wound (wTCJs) through unknown mechanisms. We found that the small GTPases Rab4 and Rab11, implicated in endosomal trafficking, are necessary for rapid wound repair. Manipulations of Rab4 and Rab11 activity reduced wound closure rates but did not have effects on tissue mechanics or myosin polarization that could explain the defect. Instead, Rab4 and Rab11 redistributed E-cadherin to accumulate at wTCJs in a process necessary for collective cell movement. Together, our results show that adherens junction reinforcement via endosomal recycling is a key step of coordinated cell migration that controls the rate of wound healing independent of cytoskeletal remodeling.

**Summary Statement:** During embryonic wound repair, Rab4 and Rab11 GTPases transport E-cadherin to reinforce specialized cell adhesions at the wound edge, which are essential to drive rapid collective cell migration.

## Introduction

Collective cell migration, or the coordinated movement of a group of cells, is a fundamental process underlying development, disease, and tissue repair (Friedl and Gilmour, 2009; Stock and Pauli, 2021). Multiple developmental processes across species rely on coordinated cell movements, including *Caenorhabditis elegans* ventral enclosure (Williams-Masson et al., 1997), neural crest cell migration in Xenopus (Shellard et al., 2018), and eyelid closure in the mouse (Heller et al., 2014). The same underlying mechanisms contribute to cancer cell invasion and metastasis (Friedl and Wolf, 2003; Zhang and Weinberg, 2018). Collective cell migration also drives wound repair in the embryonic epidermis: embryos across species are able to heal wounds rapidly, without inflammation or scarring, through migration of cells into the wounded region (Martin and Lewis, 1992; McCluskey and Martin, 1995; Davidson et al., 2002; Wood et al., 2002). Thus, understanding the molecular mechanisms that promote rapid cohesive cell movement would offer insight into numerous physiological and pathophysiological processes.

To initiate and sustain migration during wound healing, cells must rearrange their cytoskeleton and adhesions (Hunter and Fernandez-Gonzalez, 2017; Rothenberg and Fernandez-Gonzalez, 2019). Shortly following wounding, actin and the motor protein non-muscle myosin II are polarized to the leading edge of the migrating cells, resulting in the assembly of a contractile actomyosin cable around the wound edge (Martin and Lewis, 1992; Brock et al., 1996; Wood et al., 2002; Abreu-Blanco et al., 2012). At the same time, adherens junctions linking neighboring cells are also remodeled. Adherens junction components, including E-cadherin, α-catenin, β-catenin, and Canoe are endocytosed from the wound edge and accumulate at former tricellular junctions along the wound perimeter (wound TCJs, wTCJs) (Brock et al., 1996; Wood et al., 2002; Zulueta-Coarasa et al., 2014; Hunter et al., 2015; Matsubayashi et al., 2015; Rothenberg et al., 2023). Although wTCJs seem to play a role in anchoring the actomyosin cable (Danjo and Gipson, 1998; Matsubayashi et al., 2015), disruption of wTCJ reinforcement negatively affects the rate of wound closure independent of myosin polarization (Rothenberg et al., 2023), suggesting that the accumulation of adherens junction components is a critical step in collective cell migration.

Delivery of adherens junction components to sites of mechanical stress is well-documented (le Duc et al., 2010; Yu and Zallen, 2020; Cavanaugh et al., 2022), though the mechanism remains unclear. E-cadherin flows along lateral cell-cell junctions to localize apically (Woichansky et al., 2016), while Canoe travels in condensates along cell-cell interfaces (Zheng et al., 2026). The highly conserved Rab family of small GTPases can also traffic E-cadherin through the endosomal recycling pathway. Rab5, marks endocytosed vesicles and early endosomes (Grant and Donaldson, 2009), and is implicated in epithelial remodeling via E-cadherin trafficking during *Drosophila* dorsal closure (Roeth et al., 2009) and epidermal wound healing (Matsubayashi et al., 2015). Rab4 mediates sorting from the early endosome for either rapid recycling or delivery to the recycling endosome (Sluijs et al., 1992; Sönnichsen et al., 2000; Grant and Donaldson, 2009). In morphogenesis of the *Drosophila* leg disc, Rab4 mediates transport of E-cadherin and controls adhesion remodeling (Madrid et al., 2015). Perhaps the most well-studied Rab GTPase in adhesion remodeling is Rab11, which marks recycling endosomes and mediates transport and exocytosis of both newly synthesized and recycled membrane proteins (Ullrich et al., 1996; Langevin et al., 2005; Grant and Donaldson, 2009; Takahashi et al., 2012). Rab11 is responsible for trafficking E-cadherin in epidermal keratinocytes (Antiguas et al., 2024) as well as in the *Drosophila* wing disc (Classen et al., 2005), early embryonic ectoderm (Roeth et al., 2009), dorsal closure (Mateus et al., 2011), and developing trachea (Shaye et al., 2008). Although Rab8 is not typically associated with E-cadherin trafficking, it does localize to recycling endosomes (Grant and Donaldson, 2009) and regulates basolateral exocytosis (Bravo-Cordero et al., 2007; Henry et al., 2008) as well as delivery of membrane during *Drosophila* embryonic cellularization (Mavor et al., 2016). However, the role of Rab GTPases in mediating the rearrangement of adherens junctions necessary for rapid embryonic wound repair has not been investigated.

## Results

### Rab4 and Rab11 GTPases have differential localization in proximity to epidermal wounds

Polarized endocytic activity removes E-cadherin from the leading edge of embryonic epidermal wounds (Hunter et al., 2015). At the same time, E-cadherin and other adherens junction proteins are delivered to former tricellular junctions along the wound edge (wTCJs) (Brock et al., 1996; Wood et al., 2002; Abreu-Blanco et al., 2012; Zulueta-Coarasa et al., 2014; Rothenberg et al., 2023). We hypothesized that E-cadherin is trafficked via the endosomal recycling network within cells at the wound margin. Thus, we investigated the roles of Rab4 and Rab11 GTPase in embryonic wound healing. To examine the localization of Rab4 and Rab11, we performed immunofluorescence staining on wounded embryos. We first wounded embryos expressing Rab4 endogenously tagged with YFP (Dunst et al., 2015), and we stained the embryos using an antibody against YFP and an antibody against *Drosophila* E-cadherin (DE-cadherin). We found that Rab4 localized in puncta of various sizes in epidermal cells, with each cell containing a few large puncta as well as a myriad of small puncta (Figure 1A). At the wound edge, DE-cadherin puncta colocalize with small Rab4-positive structures (Figure 1A’, inset 1). To quantify changes in Rab4 abundance at the wound edge, we annotated groups of cells directly adjacent to the wound edge and in the 3^rd^ row of cells from the wound edge and measured average Rab4 intensity within those regions, normalized to the mean image intensity. We found a 43% increase in Rab4 intensity in regions at the wound edge relative to regions in cells further back from the wound (*p* = 0.031, Figure 1B).

**Figure 1.**
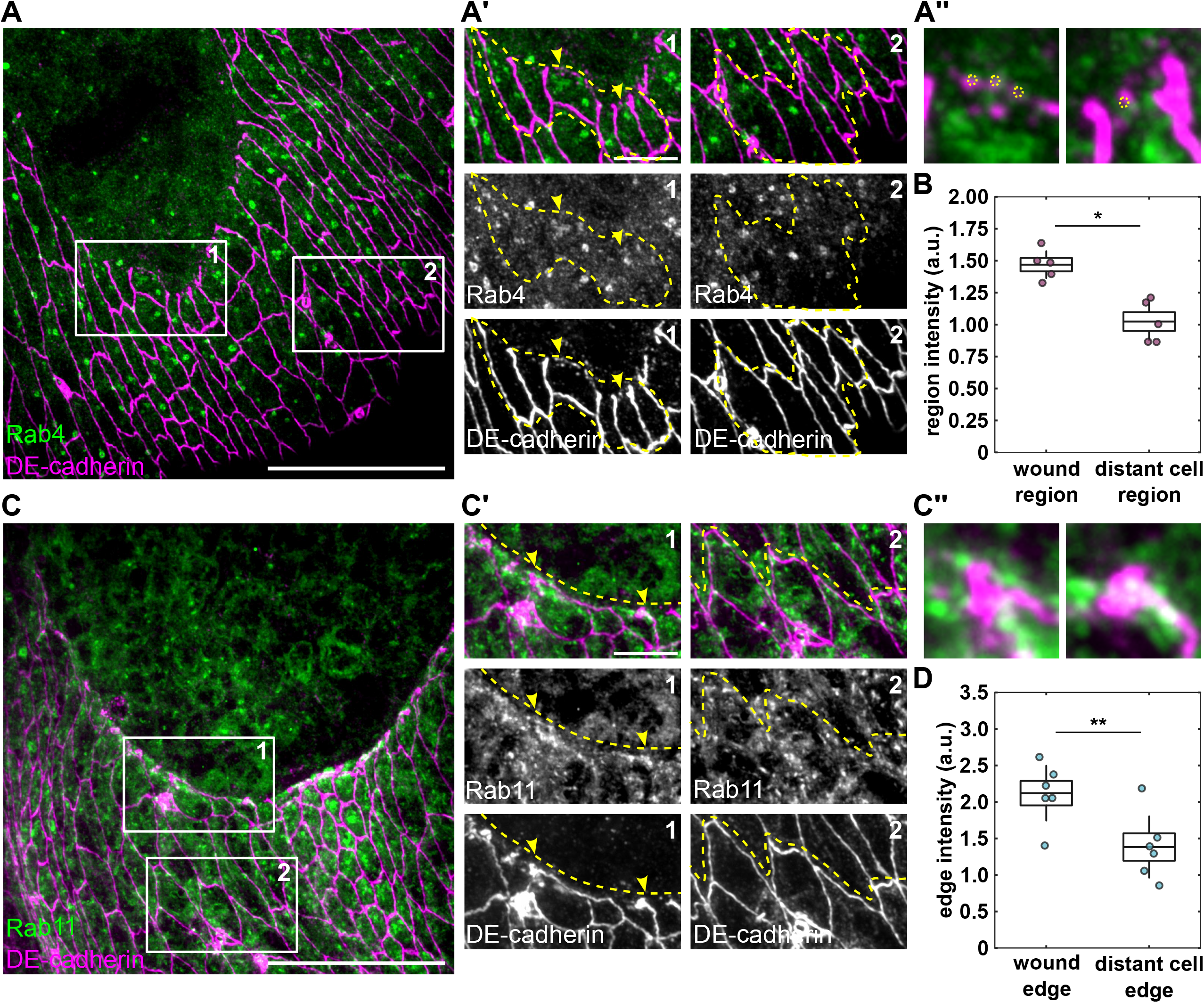
Rab4 and Rab11 GTPases show increased localization in cells surrounding embryonic epidermal wounds. **(A)** Immunofluorescence staining for Rab4 and DE-cadherin in wounded embryo expressing Rab4:YFP. (A’) shows regions indicated by bounding boxes and numbered in (A). Region 1 is adjacent to the wound edge, and region 2 is 3 rows back from the wound edge. Yellow dashed lines indicate examples of regions used for quantification. (A’’) shows regions indicated by yellow arrowheads in A’. Yellow dotted circles indicate circular structures containing both Rab4 and DE-cadherin signal **(B)** Boxplots showing mean Rab4 intensity within regions adjacent to the wound edge versus 3 rows back from the wound edge (n = 5 regions in 5 embryos). **(C)** Immunofluorescence staining for Rab11 and DE-cadherin in wounded embryo expressing Rab11:YFP. (C’) shows regions indicated by bounding boxes and numbered in (C). Region 1 is adjacent to the wound edge, and region 2 is 3 rows back from the wound edge. Yellow dashed lines indicate examples of cell edges used for quantification. (C’’) shows regions indicated by yellow arrowheads in C’. **(D)** Boxplots showing mean Rab11 intensity along cell edges adjacent to the wound edge versus 3 rows back from the wound edge (n = 6 regions in 6 embryos). (A,C) Scale bar 20 μm. Anterior left, dorsal up. (A’,C’) Scale bar 5 μm. (B,D) Error bars, s.d.; boxes, s.e.m.; \**p* < 0.05, \*\**p* < 0.01; Wilcoxon signed rank test.

Similar analysis on wounded embryos expressing Rab11:YFP (Dunst et al., 2015) revealed Rab11-positive networks within cells, rather than individual puncta (Figure 1C). Rab11 and DE-cadherin colocalized along the wound edge, particularly in proximity to wTCJs (Figure 1C’, inset 1). We found a 53% increase in Rab11 intensity along the wound edge relative to cell-cell interfaces further back from the wound (*p* = 0.031, Figure 1E). Together, our results suggest that Rab4 and Rab11 GTPases participate in the organization of the leading edge during embryonic wound healing.

DE-cadherin could also be transported towards wTCJs along cell junctions perpendicular to the wound, either by directed diffusion through the membrane or via edge-vertex flow described for the adherens junction-associated linker Canoe (Zheng et al., 2026). To test this hypothesis, we photobleached regions of bicellular junctions separating anterior-posterior neighbors (AP junctions) in either the intact epidermis or perpendicular to epidermal wounds in embryos expressing DE-cadherin endogenously tagged with GFP (Huang et al., 2009) (Figure S1A). Fluorescence recovery in AP junctions in the intact epidermis was 52 ± 8% of the initial value (Figure S1B-C), with apparently uniform fluorescence (Figure S1D). Bleached regions of AP junctions adjacent to wounds exhibited very little recovery in comparison, with a mobile fraction of 26 ± 6% (*p* = 0.03 relative to intact junctions, Figure S1B-C). We also found that bleached regions did not show any asymmetry in recovery that could indicate movement of DE-cadherin along the interface towards the wTCJ (Figure S1E). These findings support that transport along bicellular cell junctions is not a major contributor to DE-cadherin reinforcement at wTCJs.

### Active Rab4 GTPase is required for rapid wound healing by ensuring delivery of E-cadherin to former tricellular junctions

Since Rab4 puncta are more abundant in cells at the wound edge, we hypothesized that Rab4 contributes to collective cell migration and rapid wound repair. To test this hypothesis, we investigated wound closure dynamics in embryos overexpressing a dominant-negative form of Rab4 (Rab4DN) using the UAS-Gal4 system (Brand and Perrimon, 1993) and *daughterless*-Gal4 as the driver to disrupt Rab4 activity. Embryos also expressed the myosin regulatory light chain (a subunit of the myosin II motor; encoded by the *sqh* gene) tagged with mCherry (Martin et al., 2009). Embryos expressing Rab4DN took longer to close wounds than control embryos (Figure 2A).

**Figure 2.**
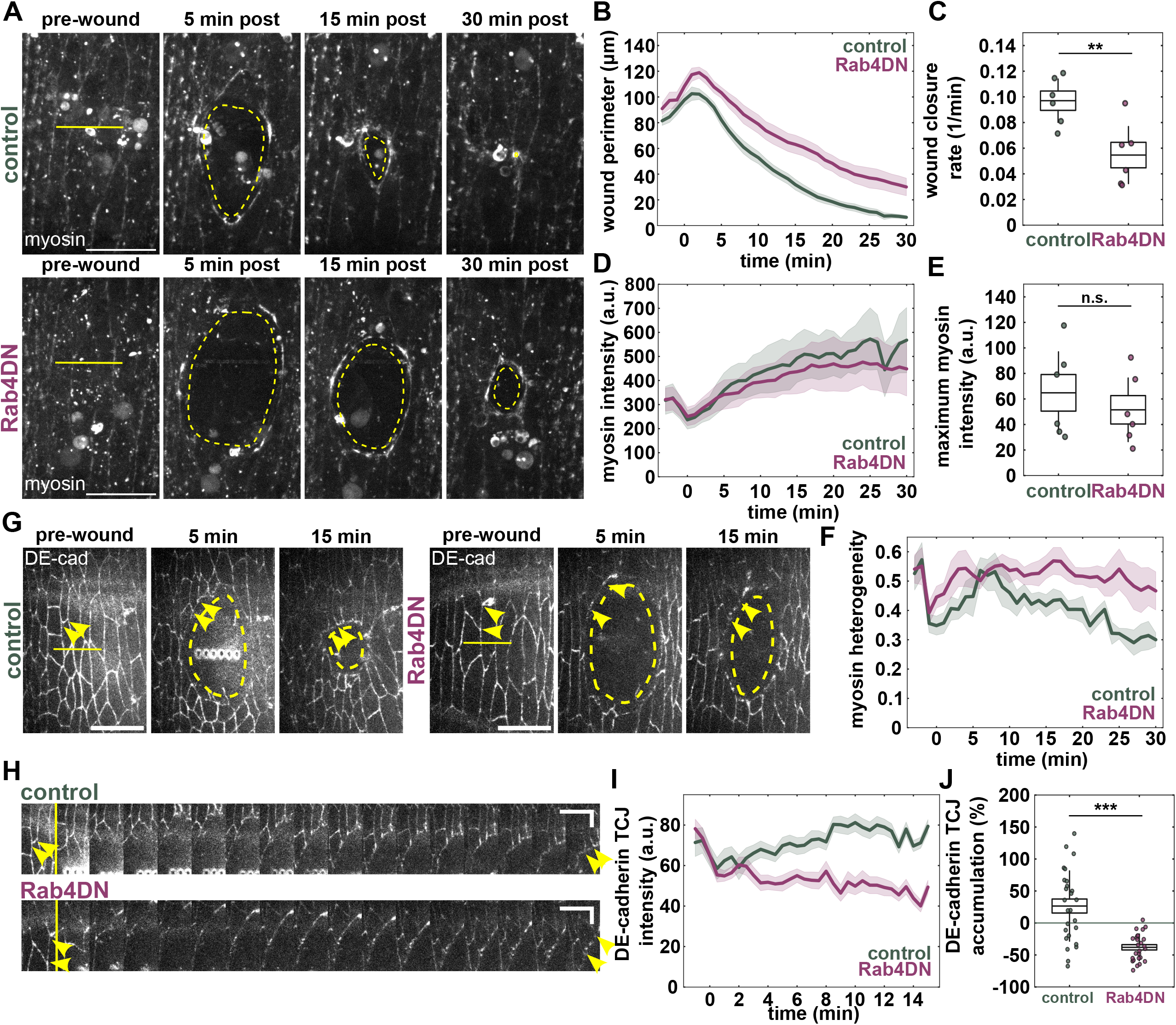
Rab4 activity is required to contribute to adhesion reinforcement that drives rapid wound healing. **(A)** Epidermal cells in wounded control and Rab4DN embryos expressing sqh:mCherry. Yellow lines denote location of wounding and yellow dashed lines indicate wound margin just interior of the wound edge. Anterior left, dorsal up. Scale bar, 15 μm. **(B-F)** Wound perimeter over time (B), boxplots of wound closure rate constant (C), average myosin intensity at the wound perimeter (D), boxplots of maximum myosin accumulation at the wound edge (E), and myosin heterogeneity over time (F) in control (n = 6, green) and Rab4DN (n = 6, magenta) embryos. **(G)** Epidermal cells in wounded control and Rab4DN embryos expressing DE-cadherin:mKate2. Anterior left, dorsal up. Scale bar, 15 μm. Yellow lines denote location of wounding, yellow dashed lines indicate wound margin, and yellow arrowheads indicate a pair of wTCJs connected by a single bicellular junction. Kymographs were generated from the indicated wTCJs. Horizontal scale bar 30 seconds, vertical scale bar 5 μm. **(H-I)** DE-cadherin intensity at wTCJs over time (H) and percent DE-cadherin intensity change 15 minutes post-wounding at wTCJs (I) in controls (n = 26 wTCJs in 4 embryos, green) and Rab4DN (n = 26 wTCJs in 4 embryos, magenta). (B,D,F,H) Shaded area represents s.e.m. (C,E,I) Error bars, s.d.; boxes, s.e.m.; \*\**p* < 0.01, \*\*\**p* < 0.001, n.s. no significance detected; Mann-Whitney test.

Quantification of the wound perimeter over time and the wound closure rate showed a 44% reduction in the rate of wound healing (*p* = 0.0087, Figure 2B-C). Thus, our results indicate that Rab4 is necessary for rapid wound closure. Overexpression of a constitutively-active form of Rab4 (Rab4CA) did not affect the rate of wound repair (Figure S2A-C), suggesting that either Rab4 is already maximally active during wound healing, or that additional activation does not affect the process.

To determine how Rab4 contributes to embryonic wound repair, we investigate how Rab4 disruption affects myosin dynamics around the wound. Myosin accumulation at the wound edge is an indicator of contractility, and defective myosin localization is often associated with slower wound healing. To measure the effects of Rab4 on myosin polarization to the wound edge, we quantified the average intensity of myosin tagged with mCherry along the wound perimeter. We found no difference in myosin intensity at the wound edge over time (Figure 2D) or in the maximal myosin accumulation during healing (Figure 2E), suggesting that Rab4 is dispensable for the delivery of myosin to the wound edge. Myosin accumulation around the wound is initially heterogenous, allowing for certain segments of the wound edge to contract while others are strained to promote additional myosin recruitment (Zulueta-Coarasa et al., 2014; Zulueta-Coarasa and Fernandez-Gonzalez, 2018). In controls, myosin heterogeneity decreases over time as all cell junctions accumulate myosin (Figure 2F). When Rab4 is disrupted, myosin heterogeneity remains increased relative to controls throughout wound closure (Figure 2F), suggesting a defect in coordination of myosin recruitment along the entire wound edge.

We previously showed that defective reinforcement of wTCJs is sufficient to slow down wound healing (Rothenberg et al., 2023). Thus, we quantified DE-cadherin intensity at wTCJs in controls and in embryos expressing Rab4DN, both co-expressing DE-cadherin endogenously tagged with 3xmKate2 (Pinheiro et al., 2017). Consistent with previous findings, DE-cadherin intensity at wTCJs increased rapidly in control embryos, within the first 15 minutes of healing (Rothenberg et al., 2023; Ly et al., 2023). In contrast, wTCJ intensity decreased in Rab4DN embryos (Figure 2G-H). We calculated the percent change in DE-cadherin intensity at wTCJs relative to intensity upon wounding as a metric of wTCJ reinforcement. Control wTCJs showed a 26 ± 11% increase in intensity at 15 minutes post-wounding, while Rab4DN wTCJs showed a 38 ± 4.1% decrease in intensity (*p* = 0.000021, Figure 2I). These results suggest that Rab4 is in part responsible for DE-cadherin delivery to wTCJs following wounding, playing an important role in orchestrating rapid migration.

### Rab11 contributes to rapid wound healing by ensuring delivery of E-cadherin to former tricellular junctions

Rab11 activity is required for rapid Xenopus embryonic wound healing, though the mechanism is unknown (Ossipova et al., 2014). To determine if Rab11 contributes to embryonic wound healing in *Drosophila*, we overexpressed a dominant-negative form of Rab11 (Rab11DN) using *daughterless*-Gal4 and measured the rate of wound closure (Figure 3A). Embryos also expressed myosin tagged with mCherry. Measurements of the wound perimeter over time and the wound closure rate showed a 50% reduction in the rate of wound healing (*p* = 0.0087, Figure 3B-C). Similarly to the Rab4DN disruption, we found no difference in the average myosin intensity at the wound edge over time (Figure 3D) or in the maximal myosin accumulation during wound closure (Figure 3E), but differences in myosin heterogeneity over time (Figure 3F). We found similar effects when we reduced *rab11* expression using an RNAi line driven with *daughterless*-Gal4 in embryos expressing myosin tagged with GFP. Wound closure in *rab11* RNAi embryos was 33% slower than in controls (*p* = 0.0022, Figure S3A-C) with no significant difference in myosin polarization to the wound edge (Figure S3D-E). Also similar to Rab4, there were no effects on wound closure resulting from overexpression of a constitutively-active form of Rab11 (Rab11CA) (Figure S2D-F). Together, these results suggest that Rab11 expression and activity are required for rapid wound healing.

**Figure 3.**
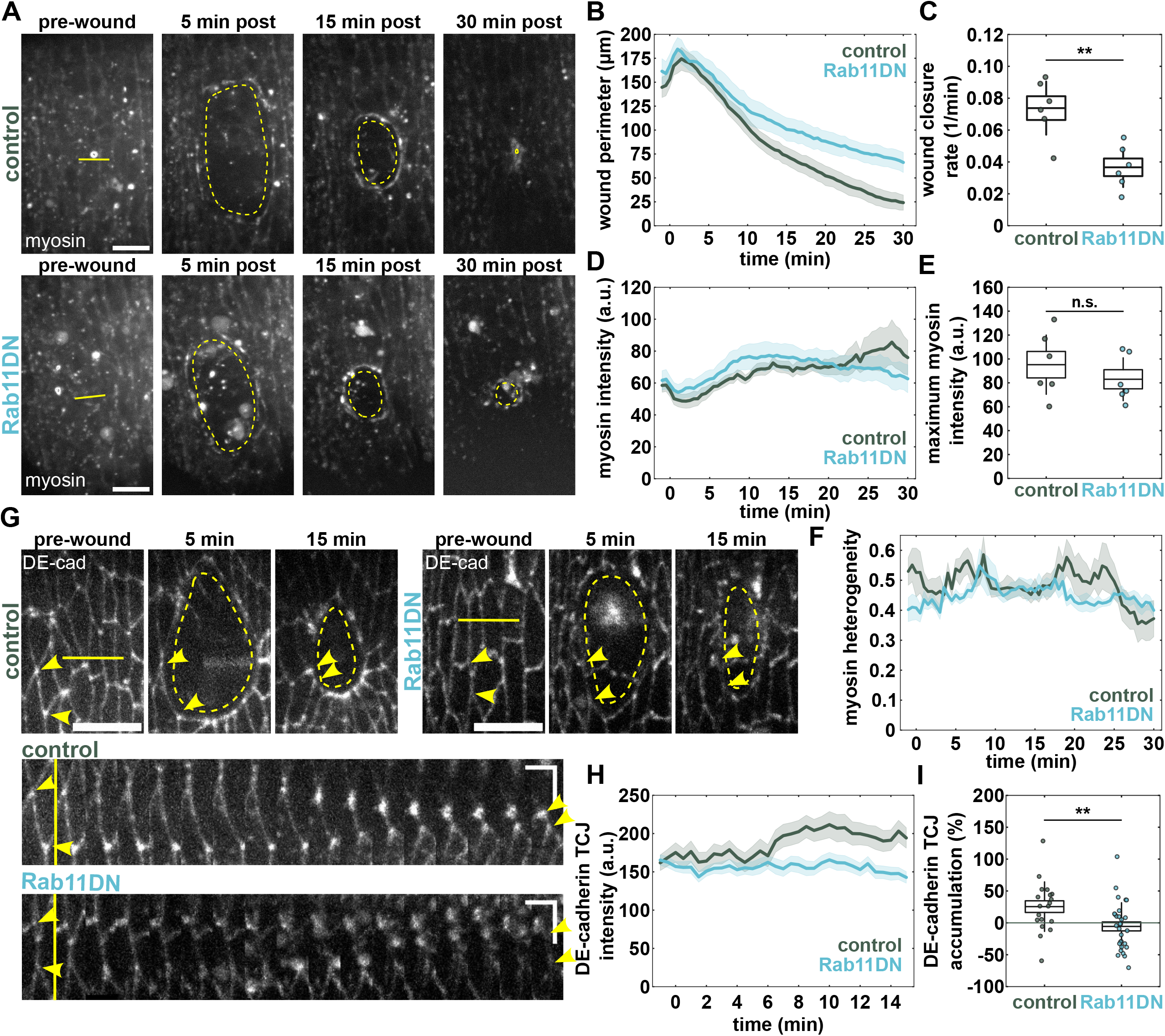
Rab11 activity is also required to contribute to adhesion reinforcement that drives rapid wound healing. **(A)** Epidermal cells in wounded control and Rab11DN embryos expressing sqh:mCherry. Yellow lines denote location of wounding and yellow dashed lines indicate wound margin just interior of the wound edge. Anterior left, dorsal up. Scale bar, 15 μm. **(B-F)** Wound perimeter over time (B), boxplots of wound closure rate constant (C), average myosin intensity at the wound perimeter (D), boxplots of maximum myosin accumulation at the wound edge (E), and myosin heterogeneity over time (F) in control (n = 6, green) and Rab11DN (n = 6, blue) embryos. **(G)** Epidermal cells in wounded control and Rab11DN embryos expressing DE-cadherin:tdTomato. Anterior left, dorsal up. Scale bar, 15 μm. Yellow lines denote location of wounding, yellow dashed lines indicate wound margin, and yellow arrowheads indicate a pair of wTCJs connected by a single bicellular junction. Kymographs were generated from the indicated wTCJs. Horizontal scale bar 30 seconds, vertical scale bar 5 μm. **(H-I)** DE-cadherin intensity at wTCJs over time (H) and percent DE-cadherin intensity change 15 minutes post-wounding at wTCJs (I) in controls (n = 19 wTCJs in 4 embryos, green) and Rab11DN (n = 29 wTCJs in 6 embryos, blue). (B,D,F,H) Shaded area represents s.e.m. (C,E,I) Error bars, s.d.; boxes, s.e.m.; \*\**p* < 0.01, n.s. no significance detected; Mann-Whitney test.

To determine if Rab11 contributes to cell adhesion remodeling during wound repair, we quantified differences in DE-cadherin accumulation to wTCJs when Rab11 was disrupted. wTCJ intensity was initially the same in controls and in Rab11DN embryos expressing DE-cadherin:tdTomato but increased steadily in controls while modestly decreasing in Rab11DN embryos (Figure 3G-H). Control wTCJs showed a 25 ± 9% increase in DE-cadherin intensity at 15 minutes post-wounding relative to the pre-wound intensity, while Rab11DN wTCJs showed a 5.6 ± 7% decrease in intensity (*p* = 0.0079, Figure 3I). *rab11* knockdown via RNAi had similar effects on wTCJs. *rab11* RNAi embryos displayed a significant 34% reduction in DE-cadherin accumulation at wTCJs relative to controls (*p* = 0.0052, Figure S3F-H). These findings establish that Rab11 plays a key role in ensuring DE-cadherin delivery to wTCJs during wound healing to promote rapid migration.

### Rab8 GTPase is not required for rapid wound healing

To ensure that the effects of Rab4 and Rab11 disruption were not due solely to perturbation of the endosomal recycling pathway, we also examined effects of disrupting Rab8 GTPase. Rab8 delivers newly synthesized proteins from the Golgi apparatus to the recycling endosome, typically for targeting to the basolateral membrane of epithelial cells for extracellular matrix ECM remodeling (Bravo-Cordero et al., 2007; Henry et al., 2008). We overexpressed a dominant-negative form of Rab8 (Rab8DN) using *daughterless*-Gal4 in embryos also expressing DE-cadherin tagged with tdTomato. Although wounds in Rab8DN were slightly larger than in controls (180 ± 7.2 μm vs. 160 ± 5.9 μm, respectively), there was no significant difference in the rates of wound closure (Figure S4A-C). Overexpression of a constitutively-active form of Rab8 (Rab8CA) also revealed no significant difference in the rate of wound closure with respect to controls (Figure S4D-F). Together, these results suggest that Rab8 does not drive wound closure, likely due to its primary role in trafficking newly synthesized proteins.

### Rab4, but not Rab11, contributes to epidermal tension

Wound healing rates can be affected by the mechanical properties of the surrounding tissue, in addition to the mechanisms that are directly driving cell migration (Tetley et al., 2019). Tissue tension and viscoelasticity can be compared across experimental groups using laser ablation of a single bicellular junction (Zulueta-Coarasa and Fernandez-Gonzalez, 2015). We performed single junction laser ablation in the epidermis of control embryos, embryos overexpressing Rab4DN, and embryos overexpressing Rab11DN (Figure 4A-B). To compare viscoelasticity between groups, the distance between the TCJs flanking the ablated junction was tracked for 1 minute following ablation (Figure 4C) and fit to a Kelvin-Voigt mechanical model to obtain relaxation time as a measurement of viscoelasticity (Figure 4D). We found no difference in relaxation time between groups (14 ± 2 sec in controls, 17 ± 3 sec in Rab4DN, and 17 ± 1 sec in Rab11DN), suggesting that Rab4 and Rab11 do not control the viscoelastic properties of the epidermis Given no detectable difference in viscoelasticity, the initial recoil velocity after ablation can be used as a proxy of tissue tension. We found a reduction in recoil velocity in Rab4DN relative to both control and Rab11DN (0.42 ± 0.039 μm/s in controls, 0.27 ± 0.041 μm/s in Rab4DN, and 0.40 ± 0.038 in Rab11DN, *p* = 0.067, Figure 4E), indicating that Rab4 maintains tissue tension in the *Drosophila* embryonic epidermis. A reduction in tissue tension should reduce resistance to cell movements, accelerating wound closure (Tetley et al., 2019). Thus, it is unlikely that the reduced rate of wound healing when Rab4 activity is disrupted is due to changes in tissue mechanics but rather defects in wTCJ reinforcement. Rab11DN does not affect tissue mechanics and shows similar phenotypes in wound healing, supporting the role that junctional rearrangements and wTCJ reinforcement play in driving rapid wound repair.

**Figure 4.**
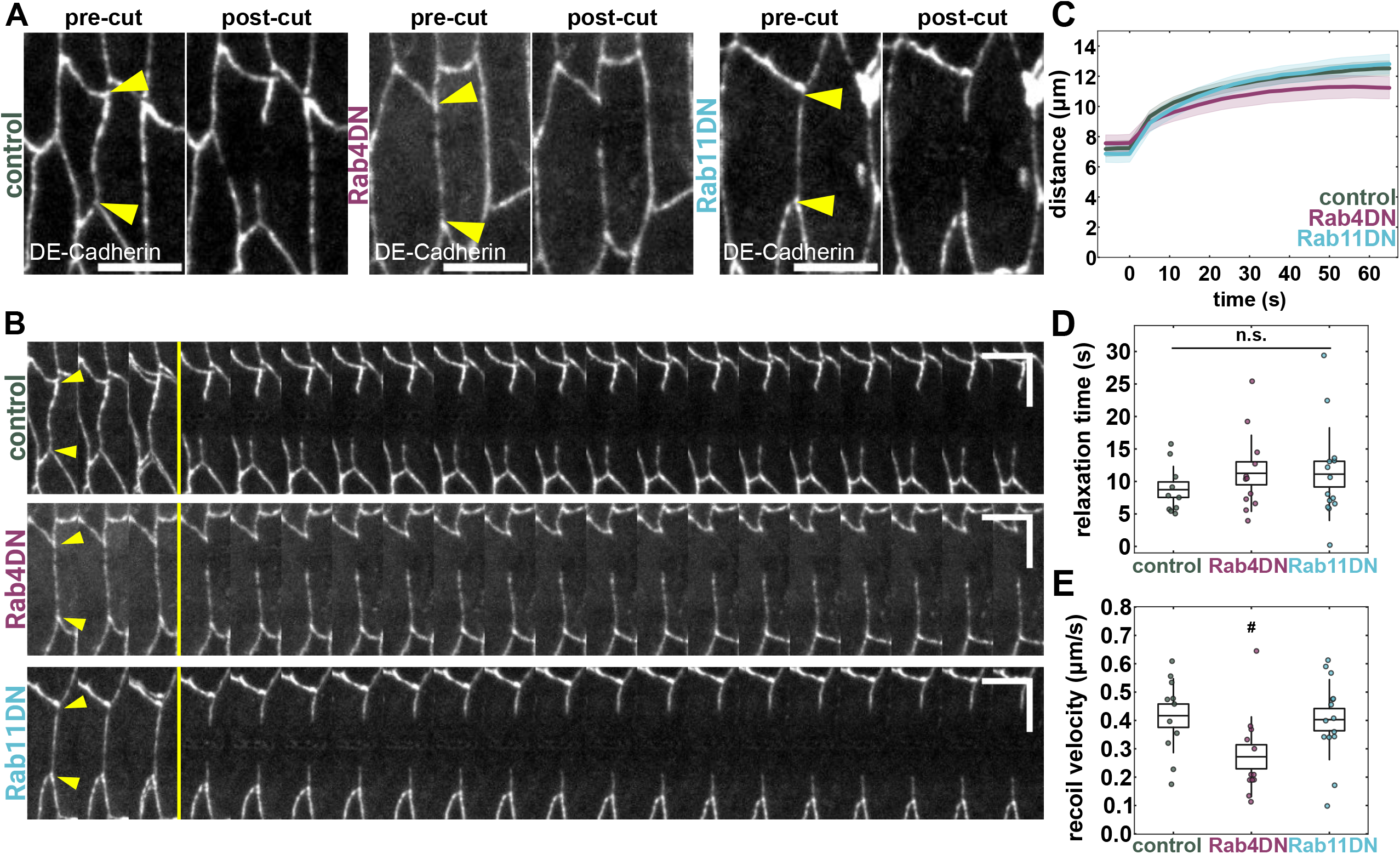
Rab4 modulates tissue tension, while Rab11 does not affect tissue mechanics. **(A)** Epidermal cells in control, Rab4DN, and Rab11DN embryos expressing DE-cadherin:GFP before and after laser ablation of a single bicellular junction. Yellow arrowheads indicate TCJs flanking cut junction. Scale bar 5 μm. **(B)** Kymographs generated from indicated junctions in (A). Yellow line denotes time of cut. Yellow arrowheads indicate TCJs flanking cut junction. Horizontal scale bar 3 seconds, vertical scale bar 5 μm **(C-E)** Distance between TCJs over time (C), boxplots of relaxation time (D), and boxplots of recoil velocity (E) for control (n = 11, green), Rab4DN (n = 12, magenta), and Rab11DN (n = 13, blue) embryos. (C) Shaded area represents s.e.m. (D,E) Error bars, s.d.; boxes, s.e.m.; ^#^*p* = 0.067; Kruskal-Wallis Test (significant), Dunn’s posthoc test with Holm correction.

### Rab4 and Rab11 work within the same pathway to drive wound closure

To understand the interplay between the functions of Rab4 and Rab11 on wound healing, we investigated the effects of simultaneously disrupting Rab4 and Rab11 activity. We compared wound closure dynamics in controls, Rab11DN embryos, and embryos expressing both Rab4DN and Rab11DN (Figure 5A). Embryos also expressed myosin tagged with mCherry. Consistent with our previous findings, we found a 54% reduction in the rate of wound healing between Rab11DN and controls (*p* = 0.028, Figure 5B-C). Simultaneous expression of Rab4DN and Rab11DN did not significantly decrease closure rate with respect to Rab11DN expression alone (Figure 5A-C). None of the disruptions showed any effect on myosin accumulation at the wound edge (Figure 5D-E), though expression of Rab11DN and Rab4DN in concert led to a sustained increase in myosin heterogeneity relative to controls (Figure 5F). DE-cadherin endogenously tagged with 3xmKate2 displayed a 38% reduction at wTCJs in Rab11DN embryos with respect to controls. (*p* = 0.00015, Figure 5F-H), and a 50% reduction in embryos co-expressing of Rab11DN along with Rab4DN (a 12% decrease relative to Rab11DN, *p* = 0.075 compared to Rab11DN alone, Figure 5F-H). Overall, our findings suggest that Rab4 may only have a small independent role in wound healing and instead may largely act upstream of Rab11 GTPase (Figure 5I).

**Figure 5.**
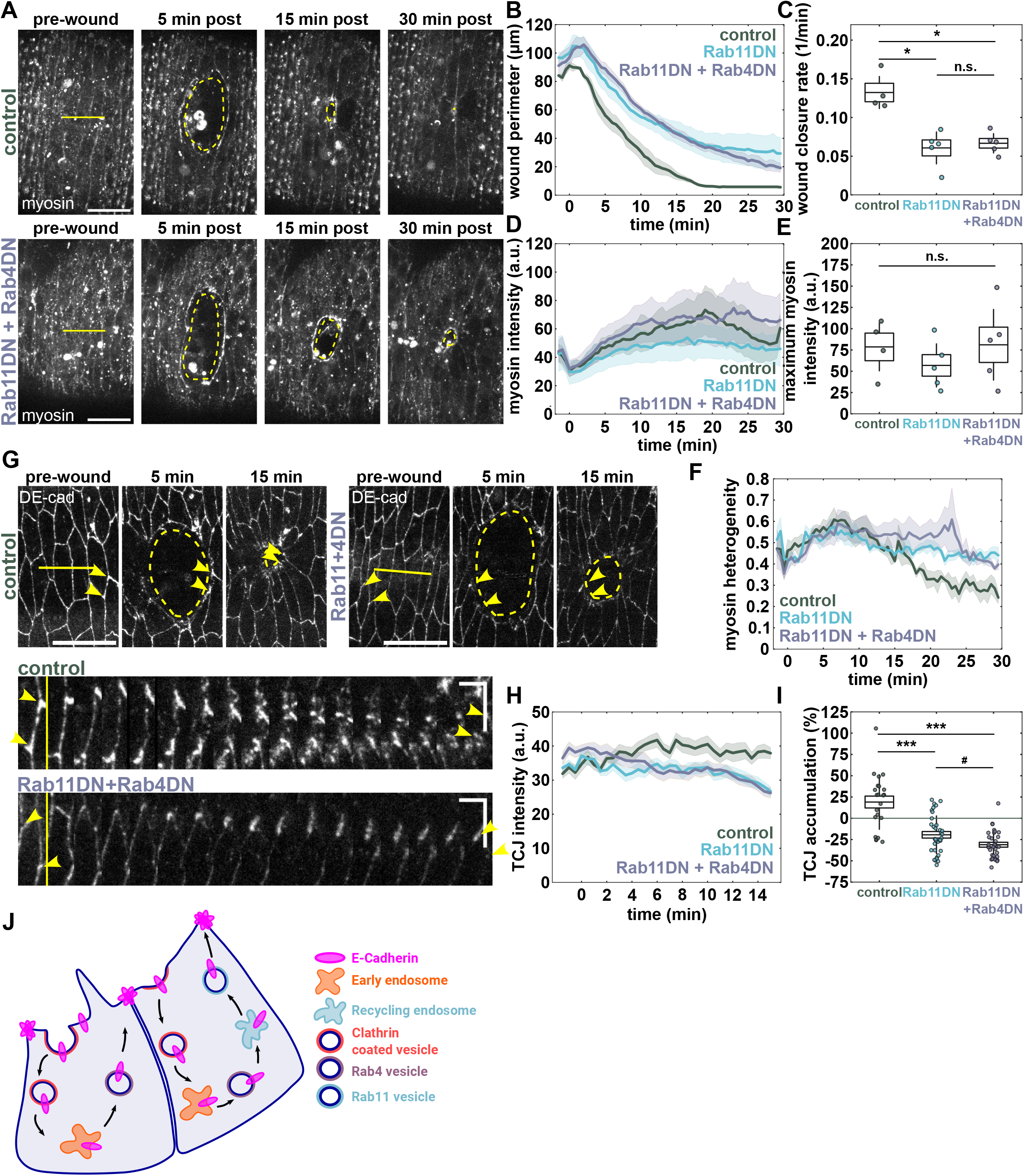
Rab4 and Rab11 seem to work together to drive wound healing. **(A)** Epidermal cells in wounded control and Rab11DN+Rab4DN embryos expressing sqh:mCherry. Yellow lines denote location of wounding and yellow dashed lines indicate wound margin just interior of the wound edge. Anterior left, dorsal up. Scale bar, 15 μm. **(B-F)** Wound perimeter over time (B), boxplots of wound closure rate constant (C), average myosin intensity at the wound perimeter (D), boxplots of maximum myosin accumulation at the wound edge (E), and myosin heterogeneity over time (F) in control (n = 4, green), Rab11DN (n = 6, blue), and Rab11DN+Rab4DN (n = 5, purple) embryos. **(G)** Epidermal cells in wounded control and Rab11DN+Rab4DN embryos expressing DE-cadherin:mKate2. Anterior left, dorsal up. Scale bar, 15 μm. Yellow lines denote location of wounding, yellow dashed lines indicate wound margin, and yellow arrowheads indicate a pair of wTCJs connected by a single bicellular junction. Kymographs were generated from the indicated wTCJs. Horizontal scale bar 60 seconds, vertical scale bar 5 μm. **(H-I)** DE-cadherin intensity at wTCJs over time (**H**) and percent DE-cadherin intensity change 15 minutes post-wounding at wTCJs (**I**) in controls (n = 22 wTCJs in 5 embryos, green), Rab11DN (n = 34 wTCJs in 5 embryos, blue), and Rab11DN+Rab4DN (n = 32 wTCJs in 5 embryos, purple). **(J)** Conceptual diagram showing model for contributions of Rab4 and Rab11 to wound healing. After E-cadherin is endocytosed from bicellular junctions, it goes to the early endosome for sorting. A small population of E-cadherin is recycled to the wTCJs directly from early endosomes by Rab4-mediated transport (left). A larger population of E-cadherin is delivered to recycling endosomes by Rab4-mediated transport, where it can be recycling to wTCJs by Rab11-mediated transport (center). (B,D,F,H) Shaded area represents s.e.m. (C,E,I) Error bars, s.d.; boxes, s.e.m.; *p< 0.05, ***p<0.001, ^#^*p* = 0.07; Kruskal-Wallis Test (significant), Dunn’s posthoc test with holm correction, n.s. no significance detected; Mann-Whitney test.

## Discussion

We demonstrate here that wTCJ reinforcement during embryonic wound healing relies on delivery of recycled E-cadherin via Rab4- and Rab11-mediated trafficking. Previous work established the requirement of clathrin-mediated endocytosis from the wound edge to enable actomyosin cable assembly (Hunter et al., 2015; Matsubayashi et al., 2015). Here, we show that the delivery of E-cadherin to wTCJs via endosomal recycling is an equally critical step in driving rapid wound closure. Reinforced wTCJs are likely anchoring the actomyosin cable and facilitating transmission and sensing of mechanical signals along the wound edge to enable rapid coordinated migration.

Rab11 has been implicated in adhesion receptor trafficking. Rab11 activity is not required for fast recycling from early endosomes but is required for slow recycling from recycling endosomes (Ren et al., 1998). Rab11-positive recycling endosomes reside between the Golgi apparatus and the cell membrane, where they receive both newly synthesized and recycled membrane proteins (Lock and Stow, 2005). Blocking protein translation globally has no effect on the wound closure process (Scepanovic et al., 2021), so it is unlikely that newly synthesized E-cadherin is delivered to recycling endosomes within the timeframe of wTCJ reinforcement (15 minutes following wounding). Together with our results that Rab8 does not play a significant role in wound closure, our findings support that the primary role of Rab11 during embryonic wound healing is delivery of recycled E-cadherin to the wTCJs to facilitate adhesion reinforcement (Figure 5I).

The exact function of Rab4 within the endosomal trafficking pathway has been difficult to determine. One suggested role of Rab4 is mediating fast recycling from the early endosome to the plasma membrane, specifically for transferrin receptor (Sluijs et al., 1992). This is further supported by delays in receptor recycling with expression of Rab4DN (Yudowski et al., 2009). (Deneka et al., 2003)There is another potential role for Rab4 in mediating delivery from early endosomes to recycling endosomes (Mohrmann and Sluijs, 1999). Rab4 can colocalize with both Rab5 and Rab11 in certain compartments, suggesting localization to both early endosomes and recycling endosomes (Sönnichsen et al., 2000). Our data support Rab4 acting in the delivery of cargo to recycling endosomes. Rab4DN and Rab11DN individually have severe wound healing phenotypes, but in combination there is no additive effect beyond a slightly enhanced defect in wTCJ reinforcement. Therefore, Rab4 may have a small role in mediating fast endosomal recycling of E-cadherin, but a larger role in transferring E-cadherin from early endosomes to recycling endosomes, where E-cadherin would be subsequently returned to the membrane by Rab11 (Figure 5I). Whether the trafficking pathway is disrupted earlier by Rab4DN expression, or later by Rab11DN expression, or both, the net result is a similar defect in wTCJ reinforcement and the rate of migration.

Rab11 may mediate trafficking of other cargo important in the wound healing process. Rab11 has been implicated in the organization of F-actin via trafficking of actin regulators, including ERM (ezrin-radixin-moesin) proteins (Desclozeaux et al., 2008; Ramel et al., 2013; Colombié et al., 2017; Plutoni et al., 2019). The role of Rab11 in actin remodeling is implicated in apicobasal polarity and lumen formation (Desclozeaux et al., 2008) as well as in collective cell migration of *Drosophila* border cell clusters (Ramel et al., 2013; Colombié et al., 2017; Plutoni et al., 2019). Actin organization also plays an important role in embryonic wound healing, with Rho GTPases controlling the formation of the actomyosin cable and actin-based protrusions that drive closure (Wood et al., 2002; Abreu-Blanco et al., 2012; Verboon and Parkhurst, 2015). Although we did not measure any changes in myosin polarization to the wound edge in response to disruptions of Rab GTPases, including Rab11, there are independent mechanisms of regulation of actin and myosin (Kobb et al., 2019). Thus, disruptions of Rab11 could affect actin organization at the wound edge without impacting myosin recruitment.

Additionally, Rab11 is known to traffic integrins to promote migration (Xu et al., 2017; Yoon et al., 2005). Integrins also play a role in driving embryonic wound closure (Campos et al., 2010; Ly et al., 2023). Therefore, future work should explore how actin dynamics and integrin localization at the wound edge are regulated by Rab11.

Fully understanding the role of Rab GTPases in collective cell migration has relevance not only for wound healing, but for development and cancer metastasis as well. Rab4, and Rab11 are implicated in morphogenetic processes that help sculpt tissues (Shaye et al., 2008; Roeth et al., 2009; Madrid et al., 2015; Chen and He, 2022), suggesting potential involvement in mechanisms underlying developmental defects. Additionally, Rab4 and Rab11 act as oncogenes in some cancers, promoting metastasis. For example, Rab4 activity contributes to melanoma progression (Barbarin and Frade, 2011), and Rab11 promotes invasion in lung cancer (Dong et al., 2017), breast cancer (Yoon et al., 2005), and cervical cancer (Xu et al., 2017). Here, we demonstrate that Rab4 and Rab11 are important for enabling effective cell coordination and migration via delivery of E-cadherin to reinforce adherens junctions. The signals that spatially target E-cadherin delivery to wTCJs remain unknown. Further investigation into the conservation of this mechanism across models of collective cell migration, as well as the upstream signals that promote targeted recycling, could yield important therapeutic targets for enhancing tissue repair or preventing cancer metastasis.

## Materials & Methods

### Fly Stocks

*Drosophila* stocks were maintained at 18°C or 25°C in plastic vials or bottles on fly food provided by a central kitchen operated by H. Lipshitz at the University of Toronto or in glass vials or bottles on fly food provided by a fly food kitchen operated by P. Clark at the University of Iowa. Embryos were collected from flies maintained at 25°C in collection cages on apple juice-agar plates with yeast paste. For immunofluorescence staining, we used endo-YFP:Rab4 (Bloomington 62542) and endo-YFP:Rab11 (Bloomington 62549) (Dunst et al., 2015). For live imaging, we used the following markers: endo-E-cadherin:GFP (Bloomington 60584) (Huang et al., 2009), endo-E-cadherin:tdTomato (Bloomington 58789) (Huang et al., 2009), endo-E-cadherin:mKate2 (Pinheiro et al., 2017), sqh>sqh:GFP (Bloomington 57145) (Royou et al., 2004), and sqh>sqh:mCherry (Bloomington 99923) (Martin et al., 2009). For disruption of proteins of interest, we used UASp>YFP:Rab4[S22N] (Bloomington 9768) (Zhang et al., 2007), UASt>Rab4 RNAi valium 20 (Bloomington 33757) (Perkins et al., 2015), UASp>YFP:Rab11[S25N](III) (Bloomington 23261) (Zhang et al., 2007), UASt>Rab11 RNAi valium10 (Bloomington 27730) (Perkins et al., 2015), UASp>YFP:Rab4[Q67L] (Bloomington 9770) (Zhang et al., 2007), and UASp>YFP:Rab11[Q70L] (III) (Bloomington 9791) (Zhang et al., 2007). UAS constructs were ubiquitously driven with *daughterless*>Gal4 (Perrin et al., 2003). For controls, we used yellow white (Morin et al., 2001) and UASt>mCherry RNAi (Bloomington 35785) (Perkins et al., 2015).

### Embryo mounting

*Drosophila* embryos were dechorionated in 50% bleach for 2 min and rinsed with water. Stage 13-14 embryos (10-12 h after egg laying) were arranged ventral side up on apple juice-agar plates using forceps. Embryos were transferred to a coverslip coated with a thin layer of heptane glue and covered with a 50/50 mixture of halocarbon oil 27 and 700.

### Time-lapse imaging and laser ablation

Time-lapse imaging and laser ablation were performed at room temperature on one of two spinning disk confocal systems.

Imaging was conducted on a Revolution XD spinning disk confocal microscope (Andor Technology) or on a Marianas Zeiss Axio Observer spinning disk confocal microscope (3i Technology). The Revolution XD confocal was equipped with a CSU-X scanner unit (Yokogawa), an iXon Ultra 897 camera (Andor Technology) or a Prime 95B camera (Teledyne Photometrics), and Metamorph software (Molecular Devices). 16-bit Z-stacks were acquired using a 60x oil-immersion lens (NA 1.35; Olympus) or a 100x oil-immersion lens (NA 1.4; Olympus) at 0.5 µm steps every 4–30 s (11–21 slices per stack). Ablations were created using a pulsed Micropoint nitrogen laser (Andor Technology) tuned to 365 nm. The Marianas confocal was equipped with a CSU-W1 SoRa scanner unit (Yokogawa), a Prime 95B camera (Teledyne Photometrics), and Slidebook software (3i). 16-bit Z stacks were acquired using a 40x oil-immersion lens (NA 1.3; Zeiss) with an additional 2.8x or 4x magnification at 0.34 μm steps every 3-60 seconds (7-19 slices per stack). Ablations were created using a 532 nm pulsed laser (3i Ablate!).

### Embryo fixation and staining

Stage 14 embryos were dechorionated, mounted on glass coverslips, and wounded. Wounded embryos were removed from the coverslips using heptane and fixed for 30 minutes in a 4% paraformaldehyde (Electron Microscopy Systems) solution in PBS. Embryos were devitellinized by hand, blocked for 1 hour in 10% BSA in PBS, and stained with primary antibodies rabbit anti-GFP (Torrey Pines Biolabs Inc) at 1:200 and rat anti-DCAD (DSHB) at 1:50 overnight at 4°C. Embryos were then incubated with secondary antibodies anti-rabbit Alexa Fluor Plus 488 (Invitrogen) at 1:500, anti-rat Alexa Fluor Plus 555 (Invitrogen) at 1:500, and Alexa Fluor Plus 647 phalloidin (Invitrogen) at 1:500 and incubated for 1 hour at room temperature. Stained embryos were mounted in VECTASHIELD Antifade mounting medium (Vector Laboratories) for subsequent imaging on the Marianas Zeiss Axio Observer spinning disk confocal microscopy described above.

### Laser ablation assays

Wounds were created on the ventrolateral embryonic epidermis. Each embryo was wounded only once. Lines with lengths from 8-14 μm were drawn across 3-4 cells for targeted ablation. The laser was set to fire along the line for 3-10 repetitions to sever cell-cell interfaces and induce cell death in targeted cells.

For epidermal tension and viscoelasticity measurements, single junction cuts were made on the ventrolateral embryonic epidermis. Each embryo was targeted only once. A 3.2 μm line was placed perpendicular to a single cell-cell interface for targeted ablation. The laser was set to fire along the line for 10 repetitions.

### Junctional E-cadherin photobleaching assay

We used a FRAPPA system (Andor) and a 488-nm laser for junctional photobleaching assays. A 1.1 μm x 1.1 μm (7 x 7 pixel) region along an anterior-posterior junction was photobleached with a laser power of 50, dwell time of 10 ms/pixel, and 3 pulses per pixel. Wounded embryos were allowed to heal for 3 minutes prior to photobleaching.

One z-stack was acquired prior to photobleaching. Photobleached regions were imaged immediately after photobleaching and every 3 seconds for 90 seconds.

### Quantitative image analysis

Image visualization and quantitative image analysis was performed using FIJI (Schindelin et al., 2012), PyJAMAS (Fernandez-Gonzalez et al., 2022), and custom code written in Python (Python Software Foundation, https://www.python.org/) and MATLAB (MathWorks, Inc). Background subtraction was performed either using the mode value of the image or using the mean intensity in a 100-225 pixel^2^ region of interest outside of the embryo. Images were converted from 3D to 2D using maximum intensity projections.

Maximum intensity projections were used for analysis unless otherwise indicated. Image analysis was performed using PyJAMAS. To analyze wound closure rates, we traced wounds using the semi-automated LiveWire annotation. The LiveWire annotation uses Dijkstra’s optimal path search algorithm to find and trace the brightest path of pixels between two manually selected points (Dijkstra, 1959). We quantified fluorescence at the wound edge as the mean pixel value under a 0.5 μm mask over the wound edge created by the LiveWire annotations after correction for photobleaching. Photobleaching correction was performed for each image by dividing by the mean intensity of the image and multiplying by the mean intensity of the first image in the time series. We quantified myosin heterogeneity at time *t* as:

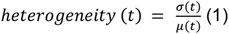

where *σ*(*t*) and *μ*(*t*) are the standard deviation and mean, respectively, of the pixel values under the wound edge annotation at time *t* (Zulueta-Coarasa and Fernandez-Gonzalez, 2018).

Time-series images to be used for wTCJ analysis was batch corrected for photobleaching. Photobleach correction curves for each image series were calculated by measuring the mean intensity of the maximum intensity projection of each time point and normalizing to the mean intensity of the first image. Curves were averaged across all control samples within an experiment and applied to all images in the experiment.

Annotations of wTCJs were manually generated by placing fiducials in PyJAMAS. wTCJs at the wound edge were tracked until 15 minutes after wounding. Average pixel intensities were measured within a 0.8 μm^2^ sized circular mask centered on the fiducial.

E-cadherin accumulation at wTCJs was measured by finding the percent change in intensity between the onset of wounding and 15 minutes following wounding.

Time-series images to be used for photobleaching analysis were first registered using PyJAMAS. Rectangular regions of interest of 7 x 7 pixels were placed at the photobleached region and in a region outside of the embryo to measure background intensity. Custom analysis in Python was used to measure normalized fluorescence intensity in the photobleached region according to established procedures (Kobb et al., 2019; Carmo et al., 2026). Mobile fraction was calculated by fitting the normalized fluorescence intensity to an exponential recovery equation and extracting the asymptotic value (Carmo et al., 2026).

To measure the recoil velocity after the single junction cut, the positions of the two TCJs connected by the junction that was cut were tracked by manually placing fiducials. We quantified the recoil velocity of TCJs after the junction cut as a proxy for mechanical tension (Zulueta-Coarasa and Fernandez-Gonzalez, 2015). We used a Kelvin-Voigt mechanical equivalent circuit, which models junctions as a spring and a dashpot configured in parallel, to estimate the tissue viscoelasticity (Zulueta-Coarasa and Fernandez-Gonzalez, 2015).

### Statistical analysis

The number of samples analyzed corresponds to the n value in all cases. When multiple samples were collected from a single embryo, as for wTCJ intensity measurements, we verified that the variance within embryos was at least as large as the variance across embryos. To measure the significance of changes across two unpaired samples, we used a non-parametric Mann-Whitney test. To measure the significance across two paired samples, we used a Wilcoxon signed-rank test. To measure the significance across three unpaired samples, we used the Kruskal-Wallis test with Dunn’s posthoc test with the Holm method of correction for multiple comparisons. In box plots, the boxes show s.e.m., the bars indicate the s.d., and the central line is the mean. In closure-rate plots, the error bars represent s.e.m. ns, not significant, \**p*<0.05, \*\**p* < 0.01, \*\*\**p* < 0.001.

## Acknowledgements

We thank Olivia Fortman, Jiarui Jiang, and Elizabeth Martin for valuable feedback on the manuscript. We thank Ana Maria Carmo for assistance with FRAP analysis. Stocks obtained from the Bloomington *Drosophila* Stock Center (NIH P40OD018537) were used in this study. We used FlyBase (release FB2026_02) to find information on stocks and gene expression (Öztürk-Çolak et al., 2024).

## Supplemental Figure Legends

**Supplemental Figure S1.**
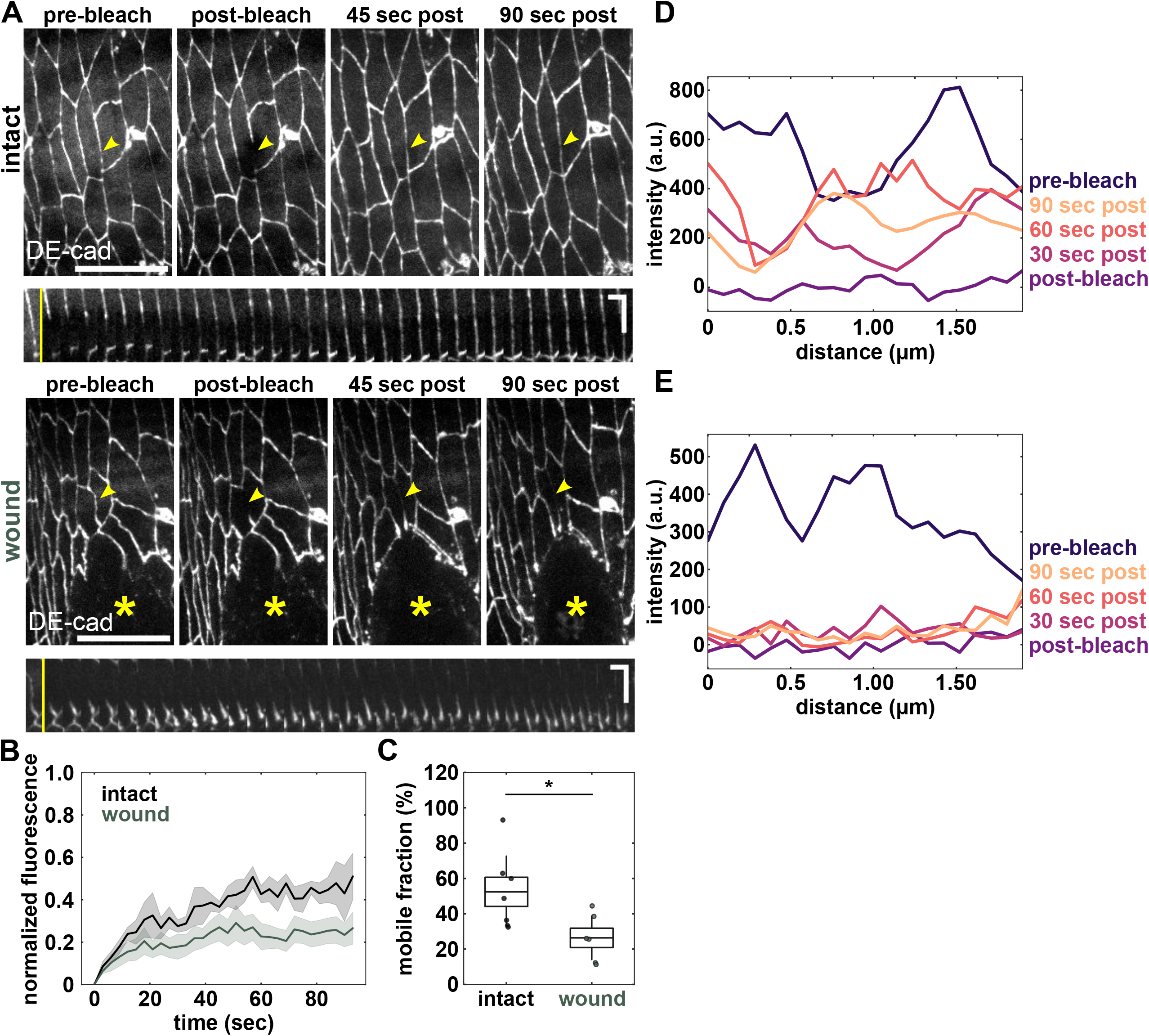
DE-cadherin does not exhibit movement along cell junctions towards wTCJs. **(A)** Photobleaching experiments in bicellular junctions in intact epidermis and adjacent to epidermal wounds in embryos expressing DE-cadherin:GFP. Yellow asterisk indicates location of wound and yellow arrowhead indicates location of bleached region. Scale bar 15 μm. Kymographs correspond to indicated bleached region with yellow line indicated time of photobleaching. Anterior left, ventral down. Horizontal scale bar 3 seconds, vertical scale bar 5 μm. **(B)** Normalized fluorescence in bleached regions showing fluorescence recovery relative to initial intensity for junctions in intact epidermis (n = 7 junctions in 7 embryos, black) and junctions adjacent to epidermal wounds (n = 6 junctions in 6 embryos, green). Shaded area represents s.e.m. **(C)** Boxplot showing mobile fraction of fluorescence recovery following fitting of normalized fluorescence in (B) with an exponential recovery function. Error bars, s.d.; boxes, s.e.m.; \**p* < 0.05; Mann-Whitney test. **(D-E)** Intensity line scans along bleached regions of junctions indicated in (A) prior to bleaching, immediately post-bleaching, and at 30 second intervals after bleaching for an intact junction (D) and a wound adjacent junction (E).

**Supplemental Figure S2.**
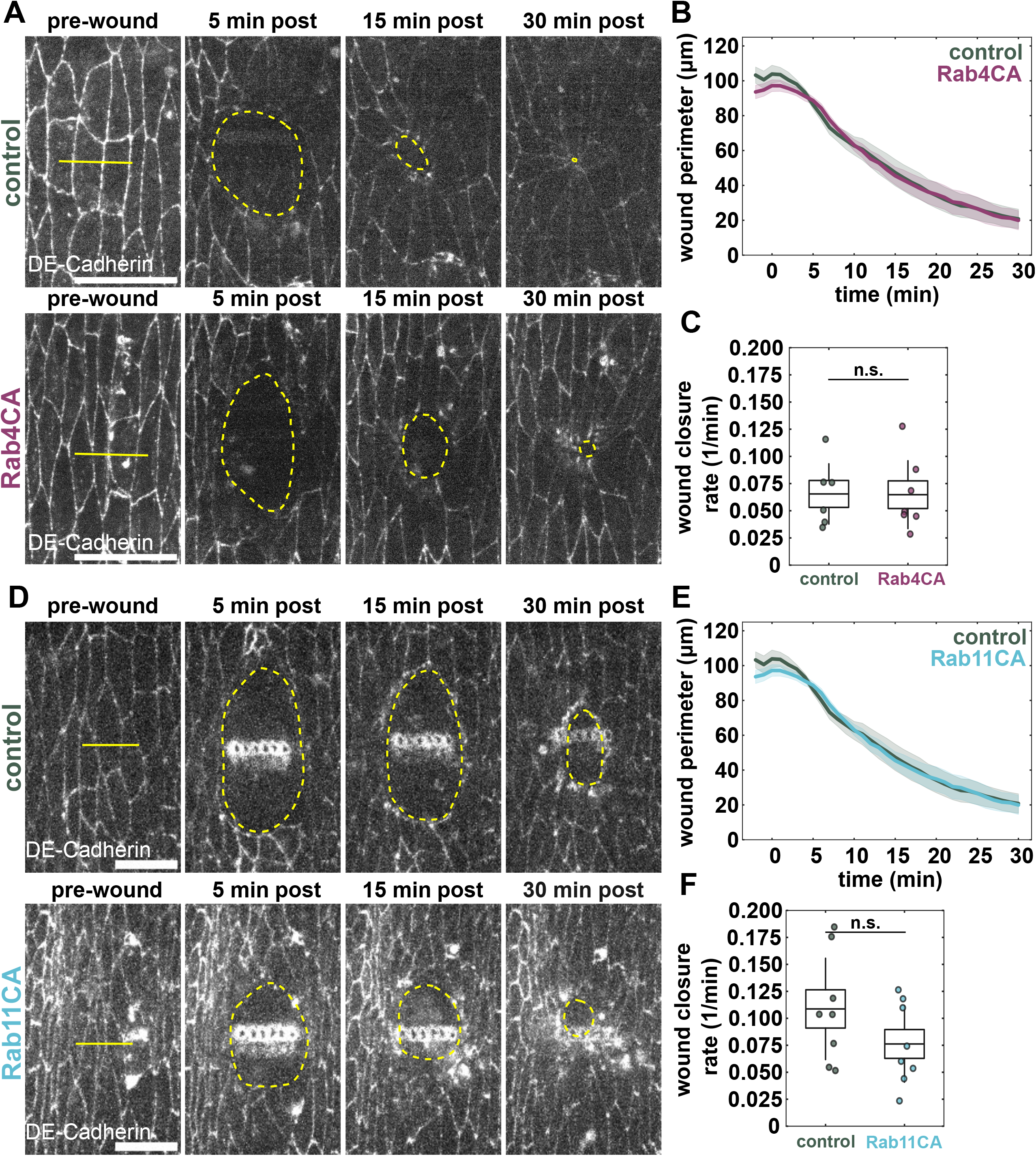
Overactivation of Rab4 and Rab11 does not affect wound closure rate. **(A)** Epidermal cells in wounded control and Rab4CA embryos expressing DE-cadherin:tdTomato. Yellow lines denote location of wounding and yellow dashed lines indicate wound margin just interior of the wound edge. Anterior left, dorsal up. Scale bar, 15 μm. **(B-C)** Wound perimeter over time (B), boxplots of wound closure rate constant (C) in control (n = 6, green) and Rab4CA (n = 7, magenta) embryos. **(D)** Epidermal cells in wounded control and Rab11CA embryos expressing DE-cadherin:tdTomato. Yellow lines denote location of wounding and yellow dashed lines indicate wound margin just interior of the wound edge. Anterior left, dorsal up. Scale bar, 15 μm. **(E-F)** Wound perimeter over time (E), boxplots of wound closure rate constant (F) in control (n = 8, green) and Rab4CA (n = 8, blue) embryos. (B,E) Shaded area represents s.e.m. (C,F) Error bars, s.d.; boxes, s.e.m.; n.s. no significance detected; Mann-Whitney test.

**Supplemental Figure S3.**
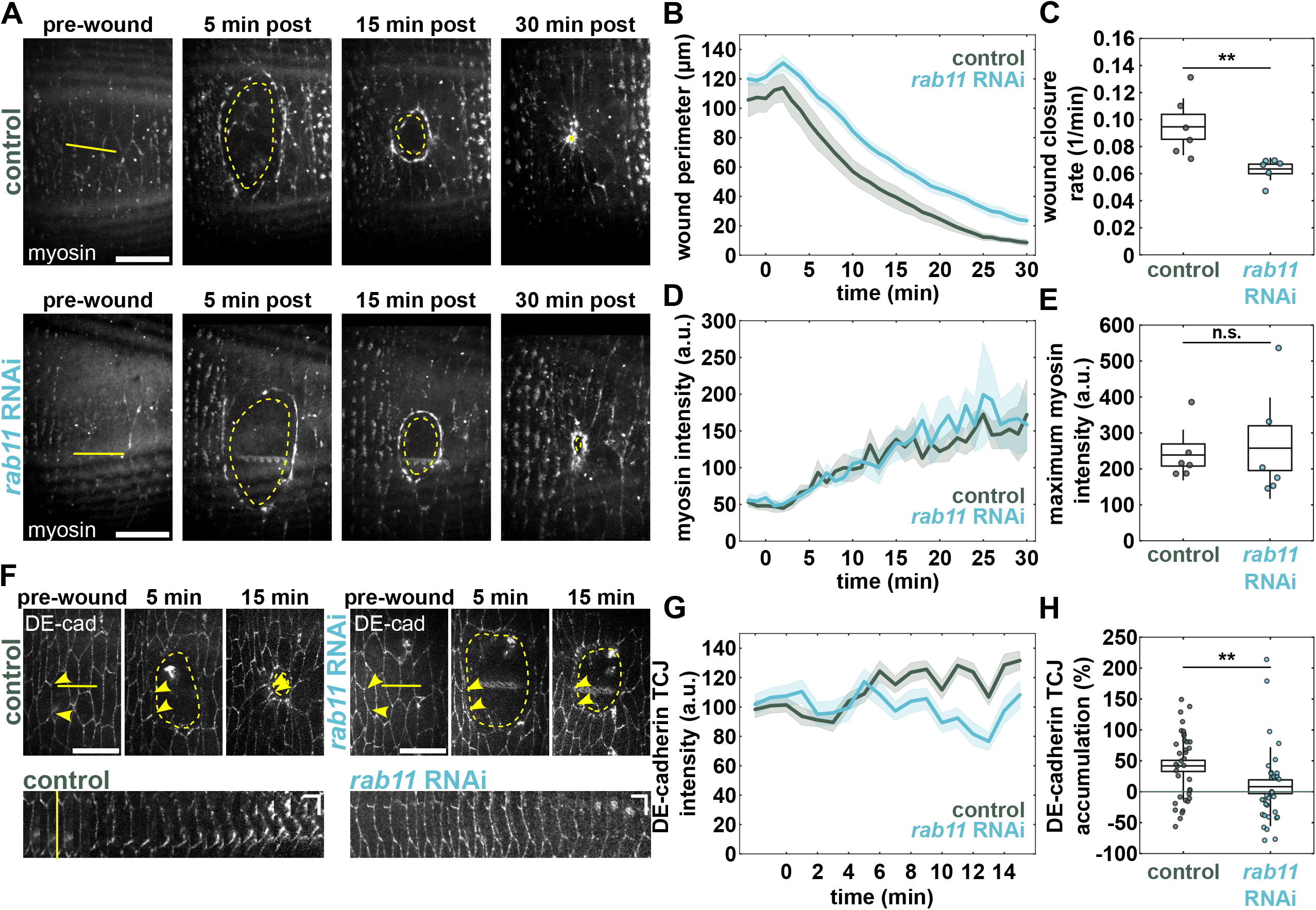
Rab11 expression is required for adhesion reinforcement that drives wound healing. **(A)** Epidermal cells in wounded control and *rab11* RNAi embryos expressing sqh:GFP. Yellow lines denote location of wounding and yellow dashed lines indicate wound margin just interior of the wound edge. Anterior left, dorsal up. Scale bar, 15 μm. **(B-E)** Wound perimeter over time (B), boxplots of wound closure rate constant (C), average myosin intensity at the wound perimeter (D), and boxplots of maximum myosin accumulation at the wound edge (E) in control (n = 6, green) and *rab4* RNAi (n = 6, blue) embryos. **(F)** Epidermal cells in wounded control and *rab11* RNAi embryos expressing DE-cadherin:tdTomato. Anterior left, dorsal up. Scale bar, 15 μm. Yellow lines denote location of wounding, yellow dashed lines indicate wound margin, and yellow arrowheads indicate a pair of wTCJs connected by a single bicellular junction. Kymographs were generated from the indicated wTCJs. Horizontal scale bar 30 seconds, vertical scale bar 5 μm. **(G-H)** DE-cadherin intensity at wTCJs over time (G) and percent DE-cadherin intensity change 15 minutes post-wounding at wTCJs (H) in controls (n = 37 wTCJs in 4 embryos, green) and *rab11* RNAi (n = 33 wTCJs in 5 embryos, blue). (B,D,G) Shaded area represents s.e.m. (C,E,H) Error bars, s.d.; boxes, s.e.m.; \*\**p* < 0.01, n.s. no significance detected; Mann-Whitney test.

**Supplemental Figure S4.**
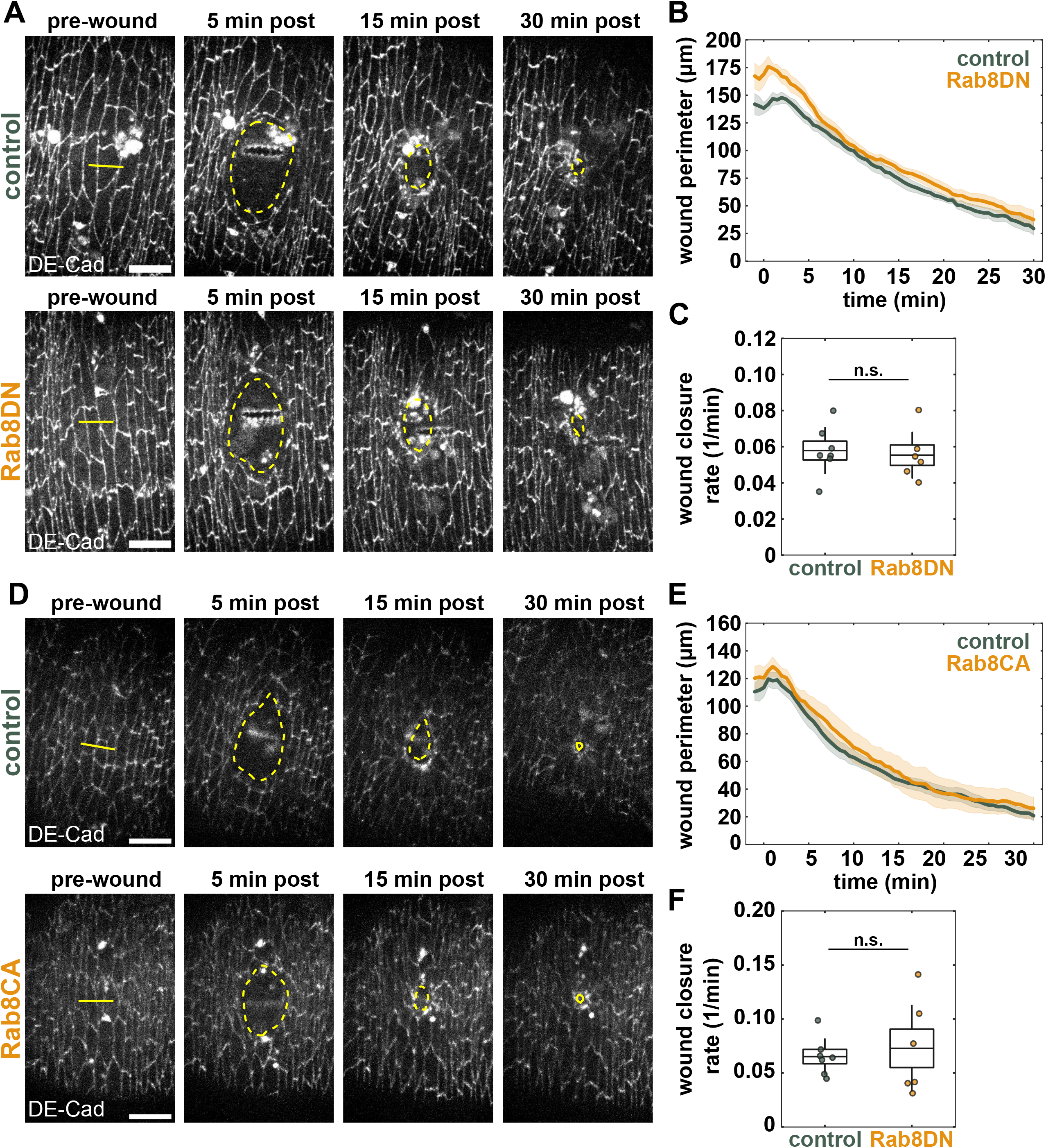
Rab8 does not play a role in epidermal wound closure. **(A)** Epidermal cells in wounded control and Rab8DN embryos expressing DE-cadherin:tdTomato. Yellow lines denote location of wounding and yellow dashed lines indicate wound margin just interior of the wound edge. Anterior left, dorsal up. Scale bar, 15 μm. **(B-C)** Wound perimeter over time (B), boxplots of wound closure rate constant (C) in control (n = 7, green) and Rab8DN (n = 6, orange) embryos. **(D)** Epidermal cells in wounded control and Rab8CA embryos expressing DE-cadherin:tdTomato. Yellow lines denote location of wounding and yellow dashed lines indicate wound margin just interior of the wound edge. Anterior left, dorsal up. Scale bar, 15 μm. **(E-F)** Wound perimeter over time (E), boxplots of wound closure rate constant (F) in control (n = 7, green) and Rab8CA (n = 6, orange) embryos. (B,E) Shaded area represents s.e.m. (C,F) Error bars, s.d.; boxes, s.e.m.

## References

Abreu-Blanco, M.T., J.M. Verboon, R. Liu, J.J. Watts, and S.M. Parkhurst. 2012. Drosophila embryos close epithelial wounds using a combination of cellular protrusions and an actomyosin purse string. J Cell Sci. 125:5984–97. doi:10.1242/jcs.109066.

Antiguas, A., M. Dunnwald, and A. Yap. 2024. A novel noncanonical function for IRF6 in the recycling of E-cadherin. Molecular Biology of the Cell. 35:ar102. doi:10.1091/mbc.E23-11-0430.

Barbarin, A., and R. Frade. 2011. Procathepsin L secretion, which triggers tumour progression, is regulated by Rab4a in human melanoma cells. Biochem J. 437:97–107. doi:10.1042/BJ20110361.

Brand, A.H., and N. Perrimon. 1993. Targeted gene expression as a means of altering cell fates and generating dominant phenotypes. Development. 118:401–15.

Bravo-Cordero, J.J., R. Marrero-Diaz, D. Megías, L. Genís, A. García-Grande, M.A. García, A.G. Arroyo, and M.C. Montoya. 2007. MT1-MMP proinvasive activity is regulated by a novel Rab8-dependent exocytic pathway. EMBO J. 26:1499– 1510. doi:10.1038/sj.emboj.7601606.

Brock, J., K. Midwinter, J. Lewis, and P. Martin. 1996. Healing of incisional wounds in the embryonic chick wing bud: characterization of the actin purse-string and demonstration of a requirement for Rho activation. J Cell Biol. 135:1097–107.

Campos, I., J.A. Geiger, A.C. Santos, V. Carlos, and A. Jacinto. 2010. Genetic Screen in Drosophila melanogaster Uncovers a Novel Set of Genes Required for Embryonic Epithelial Repair. Genetics. 184:129. doi:10.1534/genetics.109.110288.

Carmo, A.M. do, J.H. Sayo, and R. Fernandez-Gonzalez. 2026. Cofilin promotes actin turnover and flexibility to drive coordinated cell movements in vivo. 2024.12.17.628979. doi:10.1101/2024.12.17.628979.

Cavanaugh, K.E., M.F. Staddon, T.A. Chmiel, R. Harmon, S. Budnar, A.S. Yap, S. Banerjee, and M.L. Gardel. 2022. Force-dependent intercellular adhesion strengthening underlies asymmetric adherens junction contraction. Current Biology. 32:1986–2000.e5. doi:10.1016/j.cub.2022.03.024.

Chen, W., and B. He. 2022. Actomyosin activity-dependent apical targeting of Rab11 vesicles reinforces apical constriction. Journal of Cell Biology. 221:e202103069. doi:10.1083/jcb.202103069.

Classen, A.-K., K.I. Anderson, E. Marois, and S. Eaton. 2005. Hexagonal Packing of Drosophila Wing Epithelial Cells by the Planar Cell Polarity Pathway. Developmental Cell. 9:805–817. doi:10.1016/j.devcel.2005.10.016.

Colombié, N., V. Choesmel-Cadamuro, J. Series, G. Emery, X. Wang, and D. Ramel. 2017. Non-autonomous role of Cdc42 in cell-cell communication during collective migration. Developmental Biology. 423:12–18. doi:10.1016/j.ydbio.2017.01.018.

Danjo, Y., and I.K. Gipson. 1998. Actin “purse string” filaments are anchored by E-cadherin-mediated adherens junctions at the leading edge of the epithelial wound, providing coordinated cell movement. J Cell Sci. 111 ( Pt 22):3323–32.

Davidson, L.A., A.M. Ezin, and R. Keller. 2002. Embryonic wound healing by apical contraction and ingression in Xenopus laevis. Cell Motil Cytoskeleton. 53:163–76. doi:10.1002/cm.10070.

Deneka, M., M. Neeft, I. Popa, M. van Oort, H. Sprong, V. Oorschot, J. Klumperman, P. Schu, and P. van der Sluijs. 2003. Rabaptin-5α/rabaptin-4 serves as a linker between rab4 and γ1-adaptin in membrane recycling from endosomes. EMBO J. 22:2645–2657. doi:10.1093/emboj/cdg257.

Desclozeaux, M., J. Venturato, F.G. Wylie, J.G. Kay, S.R. Joseph, H.T. Le, and J.L. Stow. 2008. Active Rab11 and functional recycling endosome are required for E-cadherin trafficking and lumen formation during epithelial morphogenesis. American Journal of Physiology-Cell Physiology. 295:C545–C556. doi:10.1152/ajpcell.00097.2008.

Dijkstra, E.W. 1959. A note on two problems in connexion with graphs. Numer. Math. 1:269–271. doi:10.1007/BF01386390.

Dong, Q., L. Fu, Y. Zhao, Y. Du, Q. Li, X. Qiu, and E. Wang. 2017. Rab11a promotes proliferation and invasion through regulation of YAP in non-small cell lung cancer. Oncotarget. 8:27800–27811. doi:10.18632/oncotarget.15359.

le Duc, Q., Q. Shi, I. Blonk, A. Sonnenberg, N. Wang, D. Leckband, and J. de Rooij. 2010. Vinculin potentiates E-cadherin mechanosensing and is recruited to actin-anchored sites within adherens junctions in a myosin II–dependent manner. J Cell Biol. 189:1107–1115. doi:10.1083/jcb.201001149.

Dunst, S., T. Kazimiers, F. von Zadow, H. Jambor, A. Sagner, B. Brankatschk, A. Mahmoud, S. Spannl, P. Tomancak, S. Eaton, and M. Brankatschk. 2015. Endogenously Tagged Rab Proteins: A Resource to Study Membrane Trafficking in *Drosophila*. Developmental Cell. 33:351–365. doi:10.1016/j.devcel.2015.03.022.

Fernandez-Gonzalez, R., N. Balaghi, K. Wang, R. Hawkins, K. Rothenberg, C. McFaul, C. Schimmer, M. Ly, A.M. do Carmo, G. Scepanovic, G. Erdemci-Tandogan, and V. Castle. 2022. PyJAMAS: open-source, multimodal segmentation and analysis of microscopy images. Bioinformatics. 38:594–596. doi:10.1093/bioinformatics/btab589.

Friedl, P., and D. Gilmour. 2009. Collective cell migration in morphogenesis, regeneration and cancer. Nat Rev Mol Cell Biol. 10:445–57. doi:10.1038/nrm2720.

Friedl, P., and K. Wolf. 2003. Tumour-cell invasion and migration: diversity and escape mechanisms. Nat Rev Cancer. 3:362–74. doi:10.1038/nrc1075.

Grant, B.D., and J.G. Donaldson. 2009. Pathways and mechanisms of endocytic recycling. Nat Rev Mol Cell Biol. 10:597–608. doi:10.1038/nrm2755.

Heller, E., K.V. Kumar, S.W. Grill, and E. Fuchs. 2014. Forces Generated by Cell Intercalation Tow Epidermal Sheets in Mammalian Tissue Morphogenesis. Developmental Cell. 28:617–632. doi:10.1016/j.devcel.2014.02.011.

Henry, L., D.R. Sheff, and -Schwartz Jennifer Lippincott. 2008. Rab8 Regulates Basolateral Secretory, But Not Recycling, Traffic at the Recycling Endosome. Molecular Biology of the Cell. 19:2059–2068. doi:10.1091/mbc.e07-09-0902.

Huang, J., W. Zhou, W. Dong, A.M. Watson, and Y. Hong. 2009. From the Cover: Directed, efficient, and versatile modifications of the Drosophila genome by genomic engineering. Proc Natl Acad Sci U S A. 106:8284–9. doi:10.1073/pnas.0900641106.

Hunter, M.V., and R. Fernandez-Gonzalez. 2017. Coordinating cell movements in vivo: junctional and cytoskeletal dynamics lead the way. Current Opinion in Cell Biology. 48:54–62. doi:10.1016/j.ceb.2017.05.005.

Hunter, M.V., D.M. Lee, T.J. Harris, and R. Fernandez-Gonzalez. 2015. Polarized E-cadherin endocytosis directs actomyosin remodeling during embryonic wound repair. J Cell Biol. 210:801–16. doi:10.1083/jcb.201501076.

Kobb, A.B., K.E. Rothenberg, and R. Fernandez-Gonzalez. 2019. Actin and myosin dynamics are independent during Drosophila embryonic wound repair. MBoC. 30:2901–2912. doi:10.1091/mbc.E18-11-0703.

Kouranti, I., M. Sachse, N. Arouche, B. Goud, and A. Echard. 2006. Rab35 Regulates an Endocytic Recycling Pathway Essential for the Terminal Steps of Cytokinesis. Current Biology. 16:1719–1725. doi:10.1016/j.cub.2006.07.020.

Langevin, J., M.J. Morgan, J.-B. Sibarita, S. Aresta, M. Murthy, T. Schwarz, J. Camonis, and Y. Bellaïche. 2005. Drosophila exocyst components Sec5, Sec6, and Sec15 regulate DE-Cadherin trafficking from recycling endosomes to the plasma membrane. Dev Cell. 9:365–376. doi:10.1016/j.devcel.2005.07.013.

Lock, J.G., and J.L. Stow. 2005. Rab11 in Recycling Endosomes Regulates the Sorting and Basolateral Transport of E-Cadherin. Molecular Biology of the Cell. 16:1744– 1755. doi:10.1091/mbc.e04-10-0867.

Ly, M., C. Schimmer, R. Hawkins, K. Rothenberg, and R. Fernandez-Gonzalez. 2023. Integrin-based adhesions promote cell-cell junction and cytoskeletal remodelling to drive embryonic wound healing. Journal of Cell Science. jcs.261138. doi:10.1242/jcs.261138.

Madrid, B.H. de, L. Greenberg, and V. Hatini. 2015. RhoGAP68F controls transport of adhesion proteins in Rab4 endosomes to modulate epithelial morphogenesis of Drosophila leg discs. Developmental biology. 399:283. doi:10.1016/j.ydbio.2015.01.004.

Martin, A.C., M. Kaschube, and E.F. Wieschaus. 2009. Pulsed contractions of an actin-myosin network drive apical constriction. Nature. 457:495–9. doi:10.1038/nature07522.

Martin, P., and J. Lewis. 1992. Actin cables and epidermal movement in embryonic wound healing. Nature. 360:179–83. doi:10.1038/360179a0.

Mateus, A.M., N. Gorfinkiel, S. Schamberg, and A. Martinez Arias. 2011. Endocytic and recycling endosomes modulate cell shape changes and tissue behaviour during morphogenesis in Drosophila. PLoS One. 6:e18729. doi:10.1371/journal.pone.0018729.

Matsubayashi, Y., C. Coulson-Gilmer, and T.H. Millard. 2015. Endocytosis-dependent coordination of multiple actin regulators is required for wound healing. J Cell Biol. 210:419–33. doi:10.1083/jcb.201411037.

Mavor, L.M., H. Miao, Z. Zuo, R.M. Holly, Y. Xie, D. Loerke, and J.T. Blankenship. 2016. Rab8 directs furrow ingression and membrane addition during epithelial formation in Drosophila melanogaster. Development. 143:892–903. doi:10.1242/dev.128876.

McCluskey, J., and P. Martin. 1995. Analysis of the tissue movements of embryonic wound healing--DiI studies in the limb bud stage mouse embryo. Dev Biol. 170:102–14. doi:10.1006/dbio.1995.1199.

Mohrmann, Karin, and P. van der Sluijs. 1999. Regulation of membrane transport through the endocytic pathway by rabGTPases. Molecular Membrane Biology. 16:81–87. doi:10.1080/096876899294797.

Morin, X., R. Daneman, M. Zavortink, and W. Chia. 2001. A protein trap strategy to detect GFP-tagged proteins expressed from their endogenous loci in Drosophila. Proceedings of the National Academy of Sciences. 98:15050–15055. doi:10.1073/pnas.261408198.

Ossipova, O., K. Kim, B.B. Lake, K. Itoh, A. Ioannou, and S.Y. Sokol. 2014. Role of Rab11 in planar cell polarity and apical constriction during vertebrate neural tube closure. Nat Commun. 5:3734. doi:10.1038/ncomms4734.

Öztürk-Çolak, A., S.J. Marygold, G. Antonazzo, H. Attrill, D. Goutte-Gattat, V.K. Jenkins, B.B. Matthews, G. Millburn, G. dos Santos, C.J. Tabone, and FlyBase Consortium. 2024. FlyBase: updates to the Drosophila genes and genomes database. Genetics. 227:iyad211. doi:10.1093/genetics/iyad211.

Perkins, L.A., L. Holderbaum, R. Tao, Y. Hu, R. Sopko, K. McCall, D. Yang-Zhou, I. Flockhart, R. Binari, H.-S. Shim, A. Miller, A. Housden, M. Foos, S. Randkelv, C. Kelley, P. Namgyal, C. Villalta, L.-P. Liu, X. Jiang, Q. Huan-Huan, X. Wang, A. Fujiyama, A. Toyoda, K. Ayers, A. Blum, B. Czech, R. Neumuller, D. Yan, A. Cavallaro, K. Hibbard, D. Hall, L. Cooley, G.J. Hannon, R. Lehmann, A. Parks, S.E. Mohr, R. Ueda, S. Kondo, J.-Q. Ni, and N. Perrimon. 2015. The Transgenic RNAi Project at Harvard Medical School: Resources and Validation. Genetics. 201:843–852. doi:10.1534/genetics.115.180208.

Perrin, L., S. Bloyer, C. Ferraz, N. Agrawal, P. Sinha, and J.M. Dura. 2003. The leucine zipper motif of the Drosophila AF10 homologue can inhibit PRE-mediated repression: implications for leukemogenic activity of human MLL-AF10 fusions. Mol Cell Biol. 23:119–30. doi:10.1128/MCB.23.1.119-130.2003.

Pinheiro, D., E. Hannezo, S. Herszterg, F. Bosveld, I. Gaugue, M. Balakireva, Z. Wang, I. Cristo, S.U. Rigaud, O. Markova, and Y. Bellaïche. 2017. Transmission of cytokinesis forces via E-cadherin dilution and actomyosin flows. Nature. 545:103–107. doi:10.1038/nature22041.

Plutoni, C., S. Keil, C. Zeledon, L.E.A. Delsin, B. Decelle, P.P. Roux, S. Carréno, and G. Emery. 2019. Misshapen coordinates protrusion restriction and actomyosin contractility during collective cell migration. Nat Commun. 10:3940. doi:10.1038/s41467-019-11963-7.

Ramel, D., X. Wang, C. Laflamme, D.J. Montell, and G. Emery. 2013. Rab11 regulates cell–cell communication during collective cell movements. Nat Cell Biol. 15:317–324. doi:10.1038/ncb2681.

Ren, M., G. Xu, J. Zeng, C.D. Lemos-Chiarandini, M. Adesnik, and D.D. Sabatini. 1998. Hydrolysis of GTP on rab11 is required for the direct delivery of transferrin from the pericentriolar recycling compartment to the cell surface but not from sorting endosomes. Proceedings of the National Academy of Sciences of the United States of America. 95:6187. doi:10.1073/pnas.95.11.6187.

Roeth, J.F., J.K. Sawyer, D.A. Wilner, and M. Peifer. 2009. Rab11 Helps Maintain Apical Crumbs and Adherens Junctions in the Drosophila Embryonic Ectoderm. PLOS ONE. 4:e7634. doi:10.1371/journal.pone.0007634.

Rothenberg, K.E., Y. Chen, J.A. McDonald, and R. Fernandez-Gonzalez. 2023. Rap1 coordinates cell-cell adhesion and cytoskeletal reorganization to drive collective cell migration in vivo. Current Biology. 33:2587–2601.e5. doi:10.1016/j.cub.2023.05.009.

Rothenberg, K.E., and R. Fernandez-Gonzalez. 2019. Forceful closure: cytoskeletal networks in embryonic wound repair. Mol Biol Cell. 30:1353–1358. doi:10.1091/mbc.E18-04-0248.

Royou, A., C. Field, J.C. Sisson, W. Sullivan, and R. Karess. 2004. Reassessing the role and dynamics of nonmuscle myosin II during furrow formation in early Drosophila embryos. Mol Biol Cell. 15:838–50. doi:10.1091/mbc.e03-06-0440.

Scepanovic, G., M.V. Hunter, R. Kafri, and R. Fernandez-Gonzalez. 2021. p38-mediated cell growth and survival drive rapid embryonic wound repair. Cell Rep. 37:109874. doi:10.1016/j.celrep.2021.109874.

Schindelin, J., I. Arganda-Carreras, E. Frise, V. Kaynig, M. Longair, T. Pietzsch, S. Preibisch, C. Rueden, S. Saalfeld, B. Schmid, J.-Y. Tinevez, D.J. White, V. Hartenstein, K. Eliceiri, P. Tomancak, and A. Cardona. 2012. Fiji: an open-source platform for biological-image analysis. Nat Methods. 9:676–682. doi:10.1038/nmeth.2019.

Shaye, D.D., J. Casanova, and M. Llimargas. 2008. Modulation of intracellular trafficking regulates cell intercalation in the Drosophila trachea. Nat Cell Biol. 10:964–970. doi:10.1038/ncb1756.

Shellard, A., A. Szabó, X. Trepat, and R. Mayor. 2018. Supracellular contraction at the rear of neural crest cell groups drives collective chemotaxis. Science. 362:339–343. doi:10.1126/science.aau3301.

Sluijs, P. van der, M. Hull, P. Webster, P. Mâle, B. Goud, and I. Mellman. 1992. The small GTP-binding protein rab4 controls an early sorting event on the endocytic pathway. Cell. 70:729–740. doi:10.1016/0092-8674(92)90307-X.

Sönnichsen, B., S. De Renzis, E. Nielsen, J. Rietdorf, and M. Zerial. 2000. Distinct Membrane Domains on Endosomes in the Recycling Pathway Visualized by Multicolor Imaging of Rab4, Rab5, and Rab11. J Cell Biol. 149:901–914. doi:10.1083/jcb.149.4.901.

Stock, J., and A. Pauli. 2021. Self-organized cell migration across scales - from single cell movement to tissue formation. Development. 148. doi:10.1242/dev.191767.

Takahashi, S., K. Kubo, S. Waguri, A. Yabashi, H.-W. Shin, Y. Katoh, and K. Nakayama. 2012. Rab11 regulates exocytosis of recycling vesicles at the plasma membrane. J Cell Sci. 125:4049–4057. doi:10.1242/jcs.102913.

Tetley, R.J., M.F. Staddon, D. Heller, A. Hoppe, S. Banerjee, and Y. Mao. 2019. Tissue fluidity promotes epithelial wound healing. Nat. Phys. 15:1195–1203. doi:10.1038/s41567-019-0618-1.

Ullrich, O., S. Reinsch, S. Urbé, M. Zerial, and R.G. Parton. 1996. Rab11 regulates recycling through the pericentriolar recycling endosome. J Cell Biol. 135:913– 924. doi:10.1083/jcb.135.4.913.

Verboon, J.M., and S.M. Parkhurst. 2015. Rho family GTPase functions in Drosophila epithelial wound repair. Small GTPases. 6:28–35. doi:10.4161/21541248.2014.982415.

Williams-Masson, E.M., A.N. Malik, and J. Hardin. 1997. An actin-mediated two-step mechanism is required for ventral enclosure of the C. elegans hypodermis. Development. 124:2889–2901. doi:10.1242/dev.124.15.2889.

Woichansky, I., C.A. Beretta, N. Berns, and V. Riechmann. 2016. Three mechanisms control E-cadherin localization to the zonula adherens. Nat Commun. 7:10834. doi:10.1038/ncomms10834.

Wood, W., A. Jacinto, R. Grose, S. Woolner, J. Gale, C. Wilson, and P. Martin. 2002. Wound healing recapitulates morphogenesis in Drosophila embryos. Nat Cell Biol. 4:907–912. doi:10.1038/ncb875.

Xu, H., Y. Yuan, W. Wu, M. Zhou, Q. Jiang, L. Niu, J. Ji, N. Liu, L. Zhang, and X. Wang. 2017. Hypoxia stimulates invasion and migration of human cervical cancer cell lines HeLa/SiHa through the Rab11 trafficking of integrin αvβ3/FAK/PI3K pathway-mediated Rac1 activation. J Biosci. 42:491–499. doi:10.1007/s12038-017-9699-0.

Yoon, S.-O., S. Shin, and A.M. Mercurio. 2005. Hypoxia Stimulates Carcinoma Invasion by Stabilizing Microtubules and Promoting the Rab11 Trafficking of the α6β4 Integrin. Cancer Res. 65:2761–2769. doi:10.1158/0008-5472.CAN-04-4122.

Yu, H.H., and J.A. Zallen. 2020. Abl and Canoe/Afadin mediate mechanotransduction at tricellular junctions. Science. 370. doi:10.1126/science.aba5528.

Yudowski, G.A., M.A. Puthenveedu, A.G. Henry, Z.M. von, and S.L. Schmid. 2009. Cargo-Mediated Regulation of a Rapid Rab4-Dependent Recycling Pathway. Molecular Biology of the Cell. 20:2774–2784. doi:10.1091/mbc.e08-08-0892.

Zhang, J., K.L. Schulze, P.R. Hiesinger, K. Suyama, S. Wang, M. Fish, M. Acar, R.A. Hoskins, H.J. Bellen, and M.P. Scott. 2007. Thirty-One Flavors of Drosophila Rab Proteins. Genetics. 176:1307–1322. doi:10.1534/genetics.106.066761.

Zhang, Y., and R.A. Weinberg. 2018. Epithelial-to-mesenchymal transition in cancer: complexity and opportunities. Front. Med. 12:361–373. doi:10.1007/s11684-018-0656-6.

Zheng, Q.-H., C. Zhang, M.-X. Wang, X. Xiang, S. Zhang, Y. Wang, and H.H. Yu. 2026. Edge-vertex flow enables rapid adhesion reinforcement under tension. J Cell Biol. 225:e202601182. doi:10.1083/jcb.202601182.

Zulueta-Coarasa, T., and R. Fernandez-Gonzalez. 2015. Laser ablation to investigate cell and tissue mechanics in vivo. In Integrative Mechanobiology: Micro- and Nano-Techniques in Cell Mechanobiology. C.A. Simmons, D.-H. Kim, and Y. Sun, editors. Cambridge University Press, Cambridge. 128–147.

Zulueta-Coarasa, T., and R. Fernandez-Gonzalez. 2018. Dynamic force patterns promote collective cell movements during embryonic wound repair. Nat Phys. 14:750-+. doi:10.1038/s41567-018-0111-2.

Zulueta-Coarasa, T., M. Tamada, E.J. Lee, and R. Fernandez-Gonzalez. 2014. Automated multidimensional image analysis reveals a role for Abl in embryonic wound repair. Development. 141:2901–11. doi:10.1242/dev.106898.

